# Evolution of human pair bonds as a consequence of male-biased mating sex ratios?

**DOI:** 10.1101/2024.01.22.576327

**Authors:** Matthew C. Nitschke, Viney Kumar, Katrina E. Milliner, Kristen Hawkes, Peter S. Kim

## Abstract

Compared to our closest primate relatives, human life history involves greater longevity, which includes a distinctive postmenopausal life stage. The extension of the human lifespan (and continued fertility in old males) without lengthening female fertility directly changes the ratio of fertile males to fertile females, called the adult sex ratio (ASR). Additionally, this affects a more fine-grained ratio, the operational sex ratio (OSR), defined as the ratio of males to females currently able to conceive. Here, we construct an ODE model with minimal age structure, in which males compete for paternities using either a multiple-mating or mate-guarding strategy. Our focus is on investigating the differences of strategy choice between populations with chimpanzee-like and human-like life histories. By simulating the system, we determine the dominant strategy and its dependence on various parameter combinations. We introduce a new measure we call the lifetime paternity opportunities (LPO) of a given male strategy. The LPO directly calculates the payoffs of different male strategies and hence enables us to predict when strategies may shift. Our results show that an increase in OSR and ASR correlates well with a change in the dominant strategy from multiple mating to guarding.

## 1 Introduction

The life history and social structure of mating behavior differs significantly between humans and other primates. Although both humans and chimpanzees engage in a variety of mating strategies [36], it is only humans that form long-term preferential relationships [30]. On reaching sexual maturity, the life histories of chimps and humans diverge. A chimpanzee at maturity may expect to live a further fifteen to twenty years [13, 29], but a human hunter-gatherer at the same point may expect to live for another forty years or more [14, 15, 17]. The age of last birth in both chimps and humans is at approximately 45 years. It is very rare for a chimpanzee to reach this age in the wild or captivity, but see [39]. However, it is common for hunter-gatherer females to live in good health past 70 years of age. Thus, those females who survive this long spend half of their adult lifespan as post-fertile. This causes a male-biased sex ratio where more males compete for paternities from a smaller number of fertile females.

Recent mathematical models investigate the evolution of monogamy in humans [23, 24, 33, 34], with regard to the role of mating sex ratio and partner availability. Two such ratios appear frequently in the literature, the adult sex ratio (ASR), defined as the ratio of males to females in the fertile ages, and the operational sex ratio (OSR), which counts only the subset of adults currently capable of conception [2, 4]. Each ratio captures a different aspect of male choice; higher OSR corresponds to more competitors for each paternity opportunity, and a higher ASR to more competitors for each fertile female.

The importance of the ASR was emphasized by Fromhage and Jennions [6], with a model that explores the mechanisms behind the differences in parental investment. However, no guarding option was considered.

Schacht et al. [35] argue that a male-biased OSR only accurately predicts male strategies among a limited set of circumstances where one male can monopolize mates. A more recent model includes a mate-guarding option and confirms that the ASR is an important factor determining strategies males will tend to adopt [32]. Here, we extend this model to include the fraction that is post-fertile in females while still fertile in males and we reconsider the role of the OSR as a predictor of male mating strategies. In particular, we show that the OSR is an important factor in measuring the lifetime paternity opportunities (LPO) a male following a particular mating strategy may face. In our model, the LPO predicts the transition between regions where one strategy dominates the other.

Using a set of ordinary differential equations, we focus our attention on the chimpanzee-human relationship. For simplicity, we do not explicitly include male-male or female-female competition and disregard female choice. Although both humans and chimpanzees exhibit a wide range of mating behaviours, our focus is on two strategies: mate-guarding and multiple-mating. We explore how both longevity and birth-interval length affect these male mating strategies. Shorter birth intervals means more paternity chances as males have less time to wait for the next opportunity. This results in a lower OSR corresponding to less competition for each additional paternity. On the other hand, larger intervals require males to wait longer. This results in a higher OSR, corresponding to greater competition for each paternity. With an unbiased sex ratio, multiple-mating is the dominant strategy as waiting longer for the next opportunity takes away valuable opportunities elsewhere. However, with humanlike longevity and the resulting male-biased adult sex ratio, the incentives change. Under this scenario, there are fewer females available for each additional paternity, increasing the competition. The cumulative effect of competing for each paternity opportunity means that mate-guarding becomes more effective, even if the waiting time is relatively long. Results of this study can be used to inform more detailed future models.

## 2 Model

We construct a simple two-strategy ordinary differential equation (ODE) model, in which males either guard mates or multiply mate (possibly acquiring many paternities at nearly the same time), so there is a population *M* of multiple-mating males and a population *G* of guarding males searching for a mate. When a guarding male finds a mate, he guards her, forming a pair bond. In contrast, a multiple-mating male continues competing for new mating opportunities. For simplicity, we assume males follow pure strategies of multiple mating or mate guarding throughout their lives and sons inherit their strategies from their fathers.

The model considers ten populations: *F*, free females without dependants; *M*, multiple-mating males; *G*, unpaired guarding males; *F*_*m*_, females caring a dependent offspring of multiple-mating males; *P*_*g*_, pairs of guarding males and females with dependants; *P*, pairs of guarding males and females without dependants; *F*_*g*_, unpaired females caring for a dependent offspring of deceased or separated guarding males; *X*, post-fertile females without dependants; *X*_*m*_, post-fertile females caring for the dependent offspring of multiple-mating males; *X*_*g*_, post-fertile females caring dependent offspring of guarding males; and *Y*, post-reproductive males. We assume pair bonds can break up by death of either partner, at a constant break-up rate *χ*, or by female fertility ending. We also assume males compete for additional mates up to an age of frailty. At this point, those in pairs continue reproducing with their mate until the pair bond ends, and those without a mate retire and enter the population of post-reproductive males, *Y*. If females are still caring dependants when a pair breaks up, they will continue to care that dependant until maturity. Later, we will also consider the alternative assumption that pair bonds do not break up when female fertility ends, in which case the guarding male remains paired with the post-fertile female.

Our model is given by the ODE system

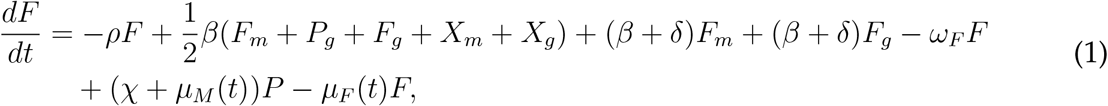

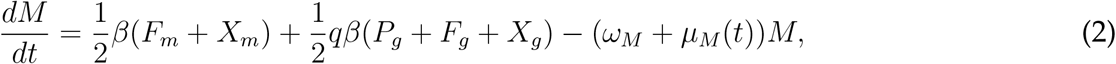

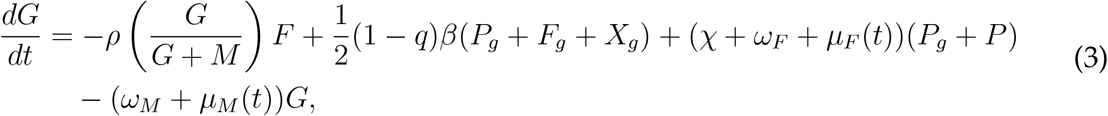

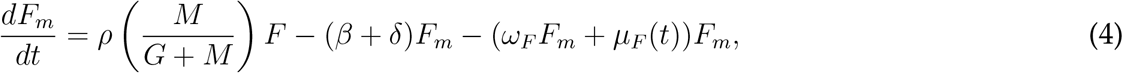

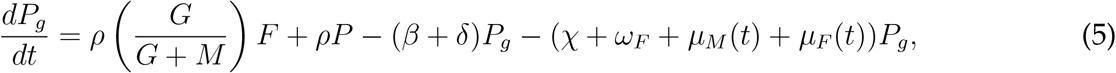

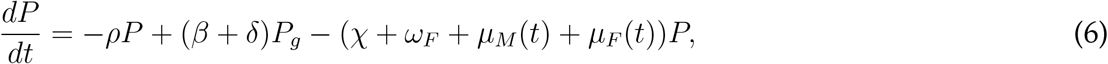

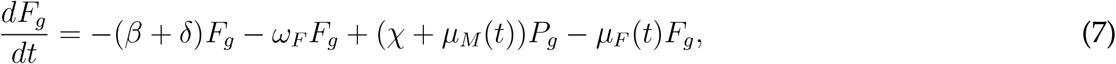

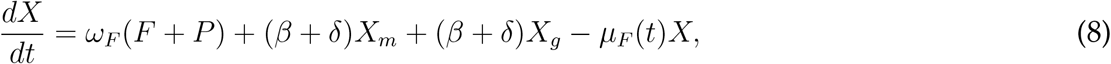

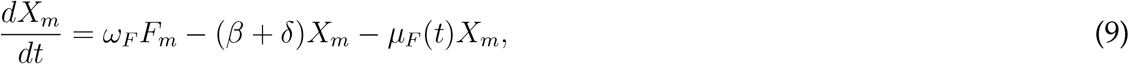

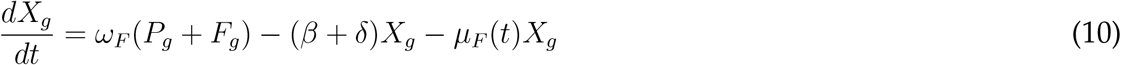

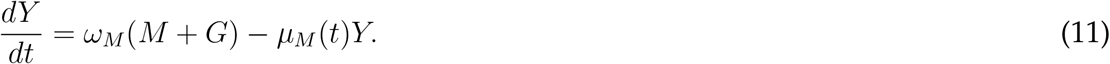

Equation (1) models the population *F* of free females without dependants. In the first term, free females conceive at rate *ρ*, causing them to enter the population *F*_*m*_ of females caring dependent offspring of multiple-mating males or *P*_*g*_ of pairs of guarding males and female mates with dependants, depending on whether they mated with multiple-mating or guarding males.

In the second term, dependants reach independence at rate *β*, allowing them to enter the adult free-female population or one of the adult male populations. All five populations caring for dependants (*F*_*m*_, *P*_*g*_, *F*_*g*_, *X*_*m*_, and *X*_*g*_) contribute maturing dependants to the adult population at rate *β*. We include a factor of 1*/*2 because we assume half the offspring are female and half are male. For simplicity, we have left out a stage of juvenile independence between dependence and adulthood.

The third term is the rate females caring dependent offspring of multiple-mating males return to the free-female population when their dependants reach independence at rate *β* or die at rate *δ*. Similarly, the fourth term is the rate unpaired females caring dependent offspring of guarding males return to the free-female population due to the independence or death of their dependants.

The fifth term is the rate free females’ fertility ends at rate *ω*_*F*_, causing them to enter the population *X* of post-fertile females without dependants. The sixth term is the rate females in pairs without dependants return to the free-female population due to their pair bonds breaking up at rate *χ* or male dying at a population-density-dependent death rate *μ*_*M*_ (*t*). The final term in (1) is the rate free females die at rate *μ*_*F*_ (*t*).

Equation (2) models the population *M* of multiple-mating males. In the first term, maturing dependants enter the adult multiple-mating-male population at rate (1*/*2)*β* from the populations *F*_*m*_ and *X*_*m*_ of fertile and post-fertile females caring offspring of multiple-mating males. As before, we include a factor of 1*/*2 because we assume half the offspring are male. The second term is the rate multiple-mating males gain dependants by stealing paternities from guarding males. Here, *q* is the level of paternity uncertainty, or the probability a dependant of a guarding male is actually the offspring of a multiple-mating male. We assume paired guarding males are unaware of paternity thefts, so the theft is only accounted in the model when dependants mature. Consequently, males may remain paired to females caring offspring of multiple-mating males. As before, *β* is the rate dependants mature, the factor of 1*/*2 accounts for the fraction of offspring who are male, and *P*_*g*_, *F*_*g*_, and *X*_*g*_ are the three populations caring dependants of guarding males. In the final term, multiple-mating males stop competing for mates due to frailty at rate *ω*_*M*_ and die at the population-density-dependent death rate *μ*_*M*_ (*t*).

Equation (3) models the population *G* of guarding males. The first term is the rate guarding males productively mate with free females. We assume productive matings occur at rate *ρF*, which is the overall conception rate of the female population. Then, for simplicity, we assume females do not favour guarding males or multiple-mating males, so the proportion of paternities going to guarding males is simply the frequency *G/*(*G* + *M*) of guarding males in the available, unpaired male population. The second term is the rate maturing dependants enter the adult guarding-male population from the populations *P*_*g*_, *F*_*g*_, and *X*_*g*_ with dependants of guarding males. It is the counterpart to the second term of (2), and the factor 1 *− q* is the probability a dependant of a guarding male is his offspring. The third term is the rate paired males return to unpaired guarding population due to pair-bond break-up at rate *χ*, female mates’ fertility ending at rate *ω*_*F*_, which we assume dissolves the pair, or female mates dying at rate *μ*_*F*_ (*t*). We assume that if females with dependants die, their dependants die, too, so unpaired males do not transition to caring any remaining dependants. In the final term, guarding males stop competing for mates due to frailty at rate *ω*_*M*_ and die at rate *μ*_*M*_ (*t*).

Equation (4) models the population *F*_*m*_ of females caring dependent offspring of multiple-mating males. The first term is the rate free females conceive with multiple-mating males.

Similarly to the first term of (3), this rate is the product of the overall female conception rate *ρF* times the frequency *M/*(*G* + *M*) of multiple-mating males. The second term is the rate dependants mature at rate *β* or die at rate *δ*, causing females in *F*_*m*_ to return to the free-female population. In the final term, females in *F*_*m*_ reach the end of their fertility at rate *ω*_*F*_, causing them to enter the population *X*_*m*_ of post-fertile females caring dependent offspring of multiple-mating males. Also, females in *F*_*m*_ die at rate *μ*_*F*_ (*t*). As in (3), we assume that if females with dependants die, their dependants die, too.

Equation (5) models the population *P*_*g*_ of pairs of guarding males and females with dependants. The first term is the rate free females conceive with guarding males, causing them to have dependants and form pairs. As in the first term of (3), this rate is the product of the overall female conception rate *ρF* times the frequency *G/*(*G* + *M*) of guarding males. The second term is the rate male-female pairs without dependants conceive at rate *ρ* and enter the population of pairs with dependants. The third term is the rate dependants mature at rate *β* or die at rate *δ*, causing pairs *P*_*g*_ with dependants to enter the population *P* without dependants. The fourth term is the rate pairs dissolve due to pair-bond break-up at rate *χ*, females in pairs ending their fertility at rate *ω*_*F*_, or either of the mates dying at rate *μ*_*F*_ (*t*) for females and *μ*_*M*_ (*t*) for males.

Equation (6) models the population *P* of pairs of guarding males and females without dependants. The first term is the rate paired females and guarding males conceive at rate *ρ*, causing them to enter the population *P*_*g*_. The second term is the rate dependants mature at rate *β* or die at rate *δ*, causing pairs in *P*_*g*_ to enter population *P*. The third term is the rate pairs dissolve due to pair-bond break-up at rate *χ*, females losing fertility at rate *ω*_*F*_, or either mate dying at rate *μ*_*F*_ (*t*) or *μ*_*M*_ (*t*).

Equation (7) models the population *F*_*g*_ of unpaired females caring dependent offspring of guarding males. This population arises when females’ mates die or pair bonds break up. The first term is the rate dependants mature at rate *β* or die at rate *δ*, causing females in *F*_*g*_ to reenter the free-female population. The second term is the rate females in *F*_*g*_ end fertility at rate *ω*_*F*_, causing them to enter the population *X*_*g*_ of post-fertile females caring dependant offspring of guarding males. The third term is the rate females in pairs with dependants enter the unpaired population *F*_*g*_ due to their pair bonds breaking up at rate *χ* or male mates dying at rate *μ*_*M*_ (*t*). The final term is the rate females in *F*_*g*_ die at rate *μ*_*F*_ (*t*). As in (4), we assume that if females with dependants die, their dependants die, too.

Equation (8) models the population *X* of post-fertile females without dependants. The first term is the rate free females in *F* and paired females in *P* end fertility at rate *ω*_*F*_, causing them to enter *X*. The second and third terms are the rates dependants mature at rate *β* or die at rate *δ*, causing post-fertile females in *X*_*m*_ and *X*_*g*_ to enter *X*. The final term is the rate females in *X* die at rate *μ*_*F*_ (*t*).

Equation (9) models the population *X*_*m*_ of post-fertile females caring dependent offspring of multiple-mating males. The first term is the rate fertile females caring dependent offspring of multiple-mating males end fertility at rate *ω*_*F*_, causing them to enter *X*_*m*_. The second term is the rate dependants mature at rate *β* or die at rate *δ*, causing females to leave *X*_*m*_ and enter the post-fertile population without dependants. The final term is the rate females in *X*_*m*_ die at rate *μ*_*F*_ (*t*).

Equation (10) models the population *X*_*g*_ of post-fertile females caring dependent offspring of guarding males. The first term is the rate paired and unpaired fertile females caring dependent offspring of guarding males lose fertility at rate *ω*_*F*_, causing them to enter *X*_*g*_. The second term is the rate dependants mature at rate *β* or die at rate *δ*, causing females to leave *X*_*g*_ and enter the post-fertile population without dependants. The final term is the rate females in *X*_*g*_ die at rate *μ*_*F*_ (*t*).

Equation (11) models the population *Y* of males who no longer compete due to frailty. The first term is the rate unpaired males in *M* and *G* become frail at rate *ω*_*M*_, causing them to enter this post-fertile population *Y*. Paired males who enter frailty remain in the pair until it ends but no longer seek additional mates afterwards. The final term is the rate males in *Y* die at rate *μ*_*M*_ (*t*).

Table 1 shows a summary of parameters and population variables. Figures 1 to 4 show model diagrams. Since the model interactions are complex, we divided them into four separate diagrams. Figure 1 shows mating interactions. Figure 2 shows transitions due to dependant maturation and death. Figure 3 shows transitions due to female fertility ending. Figure 4 shows transitions due to pair-bond break-ups and deaths of paired adults.

**Table 1:**
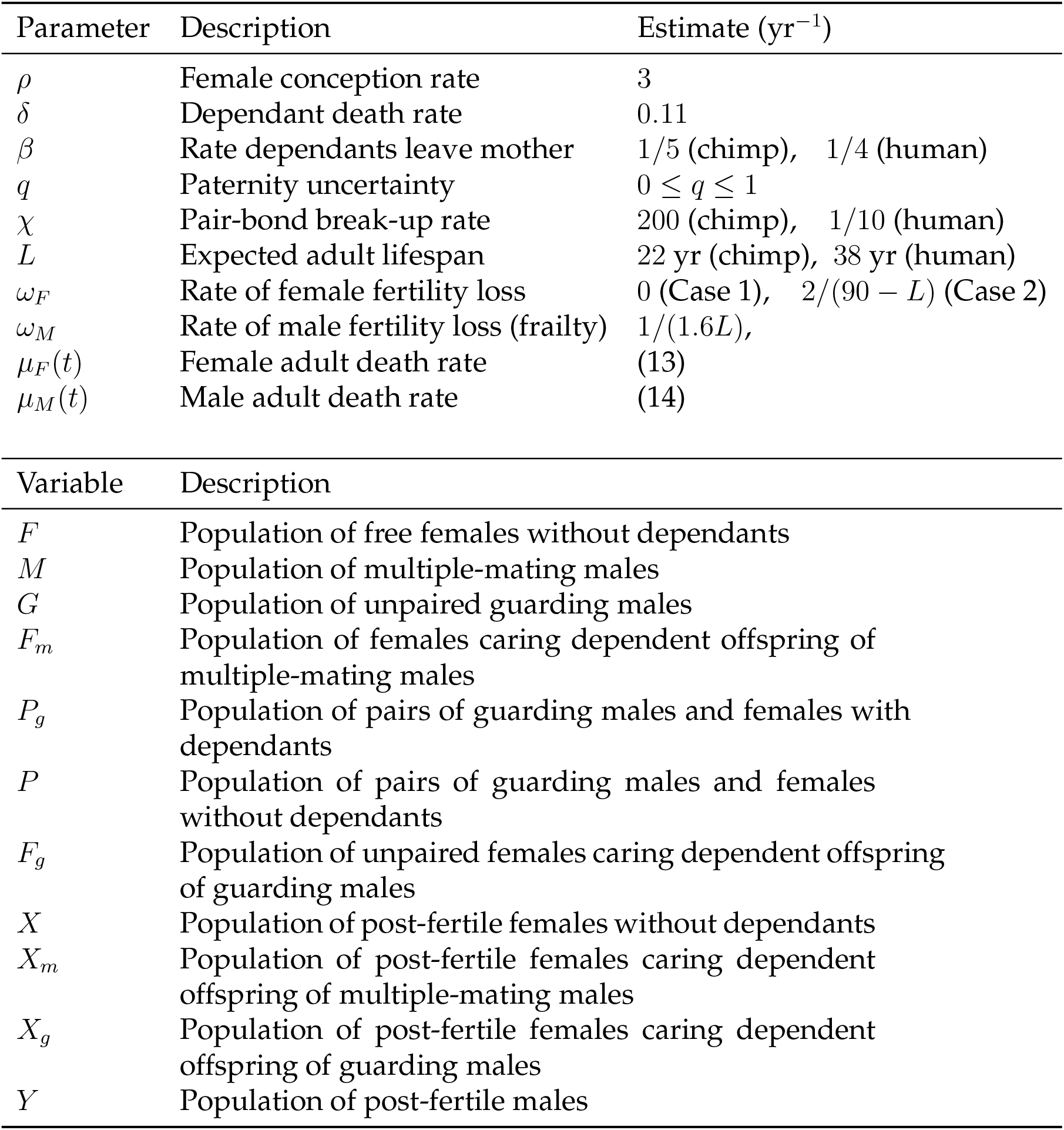
Parameters and population variables in (1)–(10). For the rate of female fertility loss, Case 1 corresponds to no female fertility not ending and hence no post-menopausal lifespan, and Case 2 corresponds to female fertility ending at age 45, which leads to an increased human post-menopausal lifespan.

| Parameter | Description | Estimate (yr<sup>-1</sup>) |
| --- | --- | --- |
| $\rho$ | Female conception rate | 3 |
| $\delta$ | Dependant death rate | 0.11 |
| $\beta$ | Rate dependants leave mother | 1/5 (chimp), 1/4 (human) |
| $q$ | Paternity uncertainty | $0 \leq q \leq 1$ |
| $\chi$ | Pair-bond break-up rate | 200 (chimp), 1/10 (human) |
| $L$ | Expected adult lifespan | 22 yr (chimp), 38 yr (human) |
| $\omega_F$ | Rate of female fertility loss | 0 (Case 1), $2/(90 - L)$ (Case 2) |
| $\omega_M$ | Rate of male fertility loss (frailty) | $1/(1.6L)$, |
| $\mu_F(t)$ | Female adult death rate | (13) |
| $\mu_M(t)$ | Male adult death rate | (14) |

**Table 1:** Parameters and population variables in (1)–(10).
| Variable | Description |
| --- | --- |
| $F$ | Population of free females without dependants |
| $M$ | Population of multiple-mating males |
| $G$ | Population of unpaired guarding males |
| $F_m$ | Population of females caring dependent offspring of multiple-mating males |
| $P_g$ | Population of pairs of guarding males and females with dependants |
| $P$ | Population of pairs of guarding males and females without dependants |
| $F_g$ | Population of unpaired females caring dependent offspring of guarding males |
| $X$ | Population of post-fertile females without dependants |
| $X_m$ | Population of post-fertile females caring dependent offspring of multiple-mating males |
| $X_g$ | Population of post-fertile females caring dependent offspring of guarding males |
| $Y$ | Population of post-fertile males |

**Figure 1:**
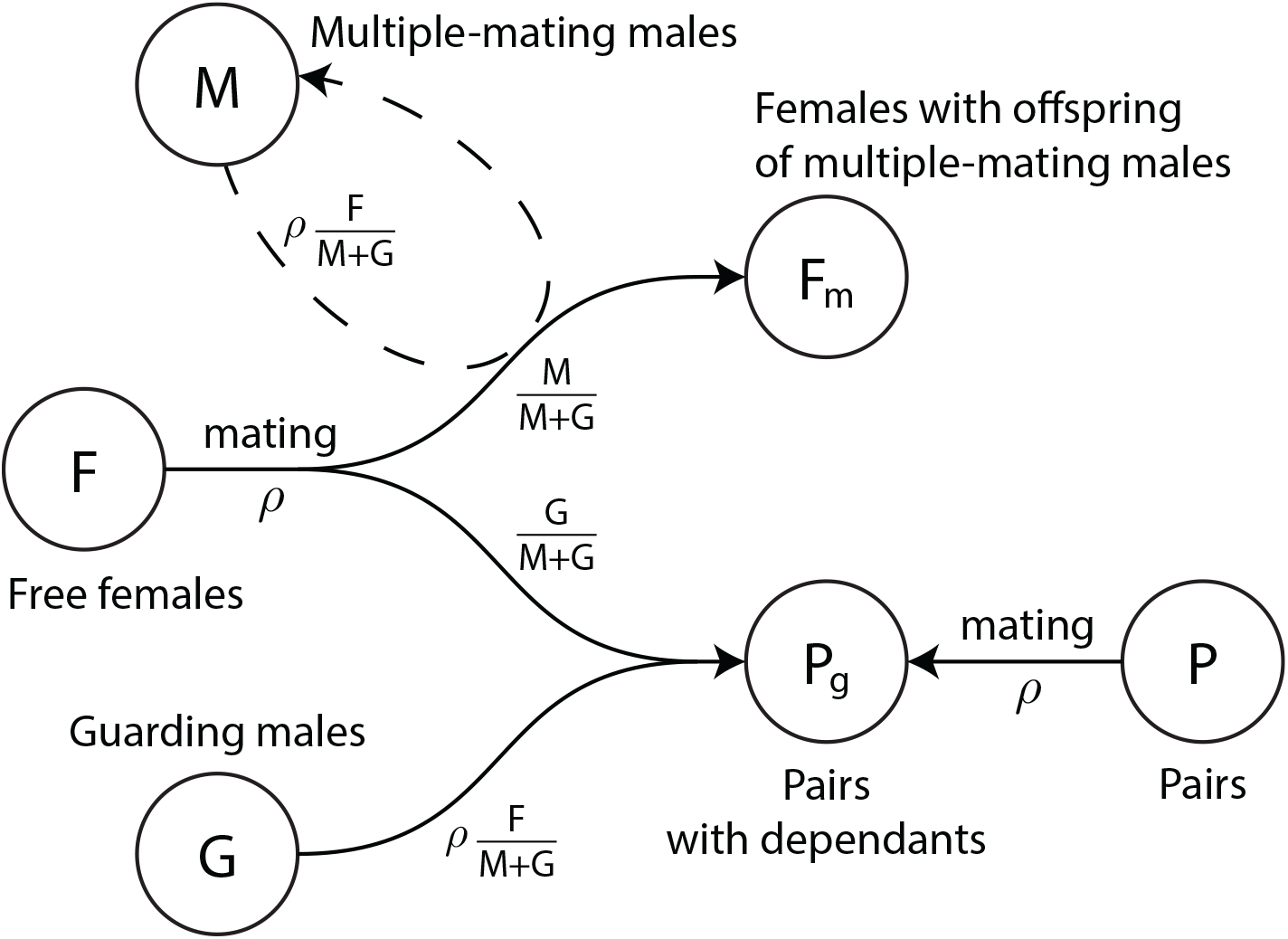
Model diagram of mating interactions. Free females, *F*, productively mate at rate *ρ*. They mate with multiple-mating males, *M*, or unpaired guarding males, *G*, with probabilities *M/*(*M* + *G*) or *G/*(*M* + *G*). If they mate with multiple-mating males, they transition to the population *F*_*m*_ of females caring offspring of multiple mating males. If they mate with guarding males, they form pairs with the guarding males and together enter the population *P*_*g*_ of pairs caring offspring of guarding males. Paternity theft can occur only after a pair is established from successful conception and is registered only after the offspring matures. (Note that females drive productive mating at rate *ρ*, so all males, *M* and *G*, mate at rate *ρF/*(*M* + *G*), which is *ρ* times the proportion of free females per unpaired male.) Pairs *P* of females and guarding males without dependants also productively mate at rate *ρ* and enter population *P*_*g*_. We include the chance that paternities from paired males can be stolen with probability *q*. In our model, this can only occur after pairs are established and the first successful conception by a guarding male. Parameters and population variables are listed in Table 1.

**Figure 2:**
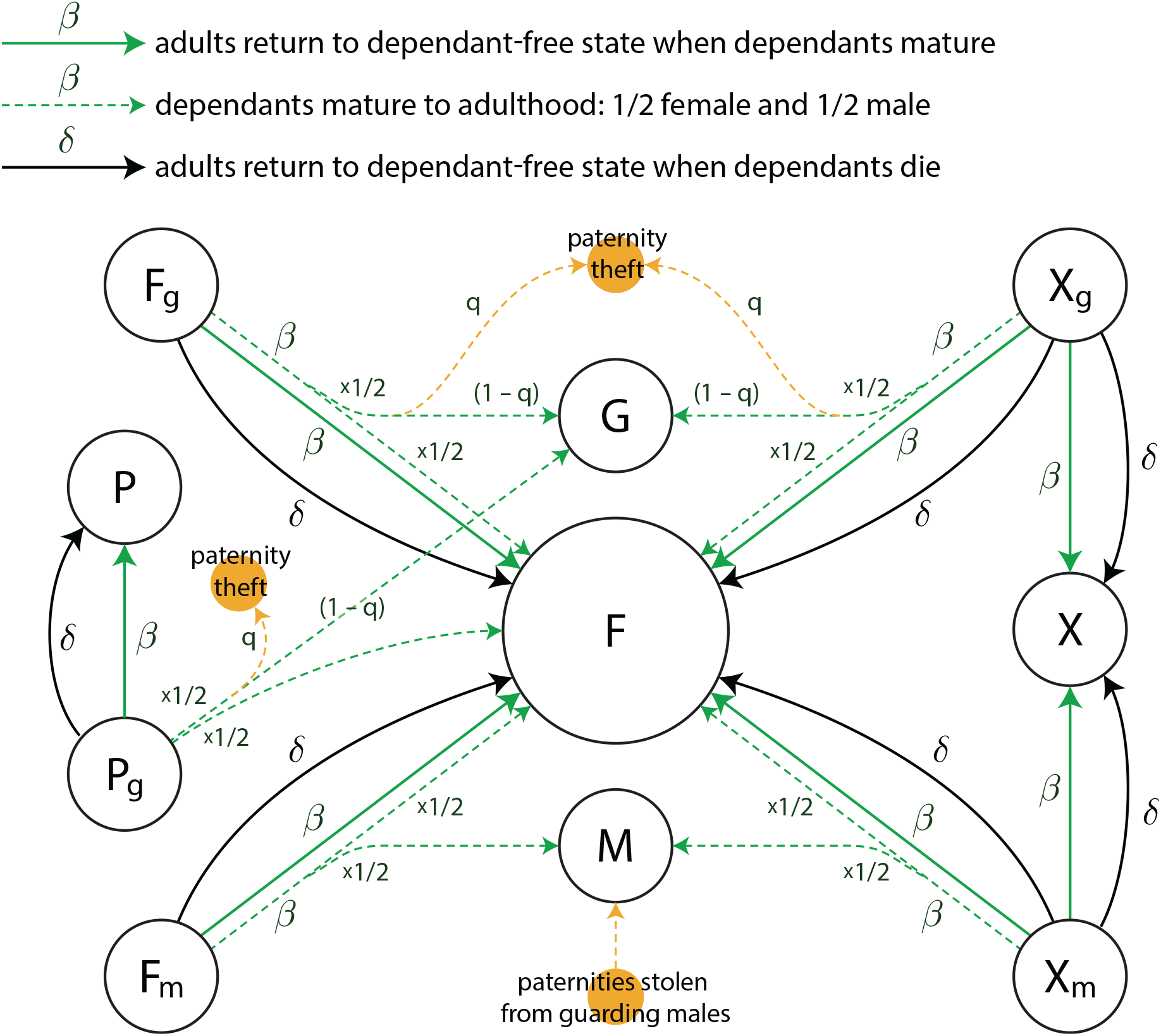
Model diagram of dependant maturation to independence and death. When dependants mature, the adult carers return to the dependant-free state, and half the maturing dependants enter the free-female population, *F*, and half enter one of the male populations, *M* or *G*. In particular, adult carers in the *P*_*g*_, *F*_*i*_, and *X*_*i*_ (*i* = *m, g*) populations return to the dependant-free populations *P, F*, and *X*, respectively. Also, maturing dependants from the *F*_*m*_ and *X*_*m*_ populations divide half and half between the *F* and *M* populations, and maturing dependants from the *F*_*g*_, *P*_*g*_, and *X*_*g*_ populations divide half and half between the *F* and *G* populations, unless the paternities are stolen by multiple-mating males with probability *q*. When dependants die, adult carers return to the dependant-free state, and there are no maturing dependants to enter the adult populations. Parameters and population variables are listed in Table 1.

**Figure 3:**
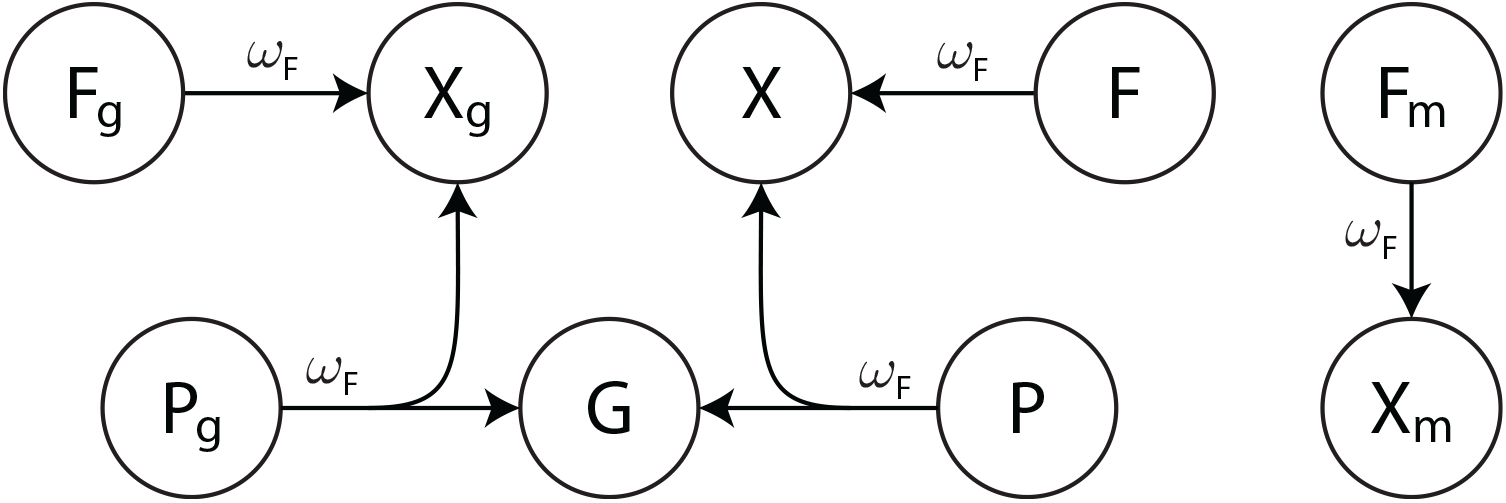
Model diagram of end of female fertility. Females end fertility at rate *ω*. When unpaired females in *F, F*_*m*_, and *F*_*g*_ end fertility, they transition to the post-fertile female populations *X, X*_*m*_, and *X*_*g*_. When females in one of the paired populations *P* or *P*_*g*_ end fertility, the pairs split up causing the males to enter the unpaired guarding-male population *G* and the females to enter *X* or *X*_*g*_ depending on whether they have dependants. Parameters and population variables are listed in Table 1.

**Figure 4:**
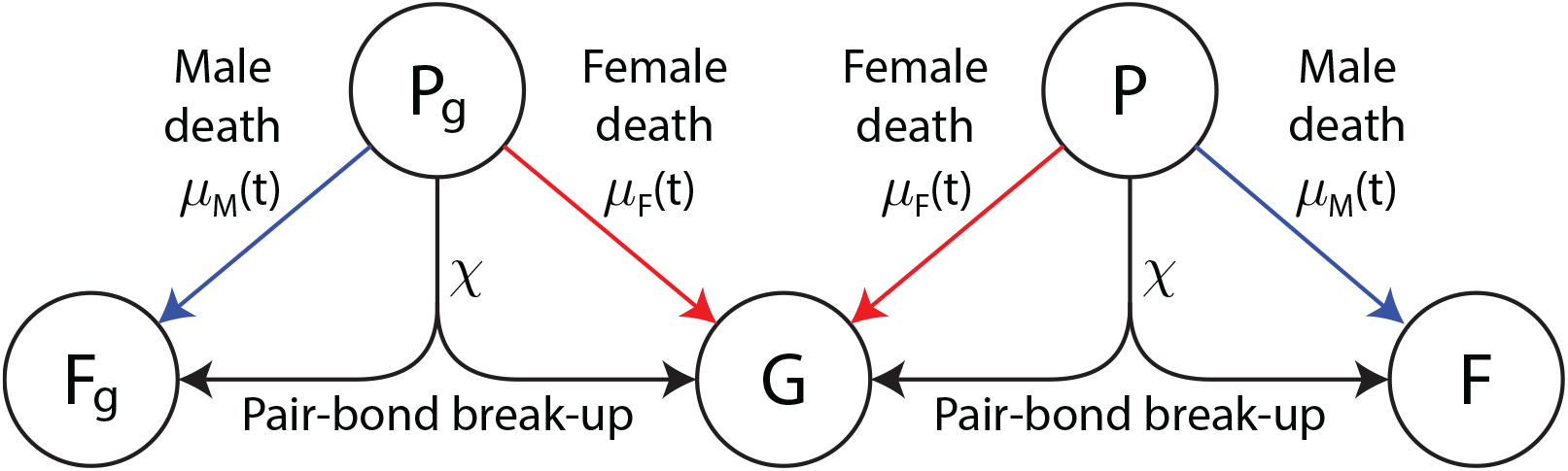
Model diagram of break-ups and deaths of paired adults. When pairs *P* or *P*_*g*_ break up at rate *χ*, the males return to the unpaired guarding population *G*, and the females return to *F* or *F*_*g*_ depending on whether they have dependants. When males in one of the paired populations *P* or *P*_*g*_ die at rate *μ*(*t*), the remaining females return to *F* or *F*_*g*_ depending on whether they have dependants. When females in one of the paired populations *P* or *P*_*g*_ die at rate *μ*(*t*), the remaining males return to *G*. We also assume all unpaired adults die at rate *μ*(*t*), but these arrows are not shown to simplify the diagram. Parameters and population variables are listed in Table 1.

Multiple-mating males will mate with available females and remain active in the mating pool after each encounter. Females who mate with these males are unavailable (time-out) until their offspring are independent or die during care. Guarding males mate and form pair bonds with their mates. Both individuals in a pair then exclusively mate with each other until the bond breaks or the female becomes infertile at a constant rate. We have included the chance that there is paternity uncertainty, which is a measure of how effective the male is at keeping an eye on his female and preventing other males from mating with her. Multiple-mating males steal paternities from paired guarding males, since they are the ones who remain with their mates. We do not distinguish between guarding males, so guarders do not steal from each other in our model.

In (1)–(11), we use a female density-dependent death rate, *μ*_*F*_ (*t*), which contains two components:

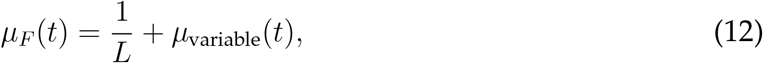

where the first term is a constant death rate given by the reciprocal of the individual’s expected adult lifespan *L* and the second term *μ*_variable_(*t*) *≥* 0 is a variable density-dependent death rate, which keeps the population constant at its initial value. In Appendix A, we derive the death rate

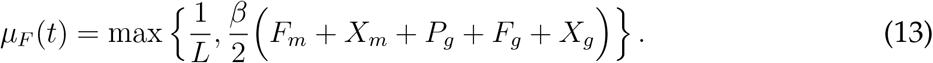

For males, we use the same density-dependent rate, but assume the baseline death rate differs by a small factor of 1.09, corresponding to a 9% higher minimum death rate. Hence, for males, we have the death rate

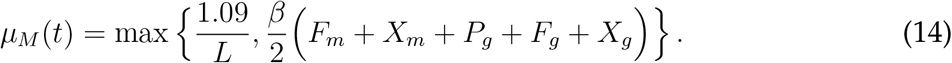

### 2.1 Parameter estimates

Conception times vary among great apes [22], but human data suggests an average of 4 months, so we estimate a female conception rate of *ρ* = 3 yr^*−*1^, corresponding to an average conception time of 1*/*3 yr, or 4 months [7]. Estimating that on average, 65% of hunter-gatherer dependants survive to age four, and wild chimpanzees show a similar survival rate [8, Table 1], we obtain a dependant death rate of δ = *−*(1*/*4) log(0.65) = 0.11 yr^*−*1^.

We estimate *β*, the rate dependants reach independence, using chimpanzee and human average interbirth intervals of 5.46 years and 3.69 years, respectively [31, Table 2.1]. Rounding these values, we assume the interbirth intervals for chimpanzees and humans are 5 and 4 years and obtain the estimates *β* = 1*/*5 and 1*/*4 yr^*−*1^, but in our simulations, we also vary this parameter more widely to get a better sense of its effect.

**Table 2:** Definition of ASR and OSR with variables in Equations 1-10. Population variables are as in Table 1.

| Ratio | Description | Definition |
| --- | --- | --- |
| $r_A$ | Adult Sex Ratio (ASR) | $\frac{M + G + P_g + P}{F + F_m + P_g + P + F_g}$ |
| $r_O$ | Operational Sex Ratio (OSR) | $\frac{M + G}{F}$ |

We include the probability *q* that fathers are uncertain of the validity of their paternities. As *q* increases, a larger proportion of youngsters cared by females paired to guarding males were sired by multiple-mating males and hence mature as multiple-mating males. In our simulations, we explore different values of this parameter, which represents guarding effectiveness.

To investigate the effect of pair-bond length on the dynamics of mating strategy, we also include the pair-bond breakup rate, *χ*. A value of *χ* = 1*/n* means that the average pair-bond lasts *n* years. Varying this parameter allows us to observe a greater range of strategies related to forming pairs.

Based on life-history regularities across primates, we assume key life-history transitions scale with respect to expected adult lifespans, *L* [1]. We assume females become sexually mature at age *L/*2. With this assumption, the age of female sexual maturity for chimpanzees and humans are 22*/*2 = 11 and 38*/*2 = 19, which fall near the ages of first birth in Robson *et al*. [31]. Then, we assume females reproduce up to age 45, which is close to the end of female fertility observed in both chimpanzees and humans [10, 38]. The average female fertile years range from sexual maturity at *L/*2 to 45, yielding a total of 45 *− L/*2 fertile years, so we estimate the average rate of female fertility loss to be the reciprocal

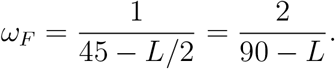

Note that this rate approximates the end of fertility, but female fecundity is constant during each female’s fertility years in our model. When a paired female’s fertility ends, her partner reenters the mating pool and competes for another female to guard. Thus, there is no decline to the benefit of guarding that corresponds with fertility loss in our model. The case when there is no post-menopausal lifespan (*ω* = 0) is discussed in Appendix 2. In this case, it is shown that multiple-mating is always the dominant strategy.

We also assume males become less competitive with age and thus retire from actively seeking additional mates during their lifetime. Note that chimpanzee and human males have generally lower fertility rates at the end of their lifetimes (age *∼* 35 in chimps and *∼* 60 in hunter-gatherers [17, 27]). This approximately scales with *L* when we observe that 60*/*38 *≈* 1.6 for humans and 35*/*22 *≈* 1.6 for chimpanzees. Thus, we estimate the average rate of male fertility loss to be the reciprocal

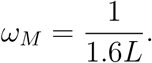

Parameter estimates are listed in Table 1.

## 3 Results

Our goal is to compare chimp- and human-like populations where males employ two possible mating strategies: multiple-mating and guarding. We do this by exploring the parameter space of the ODE model described in the previous section. More precisely, we wish to determine the dominant strategy at equilibrium corresponding to a particular set of parameters.

To provide a baseline from which to compare parameters, we start by setting the paternity uncertainty, *q*, and pair-bond break-up rate, *χ*, equal to zero. Trajectories of solutions with these assumptions are shown in Figure 5 for populations with parameters found in Table 1.

**Figure 5:**
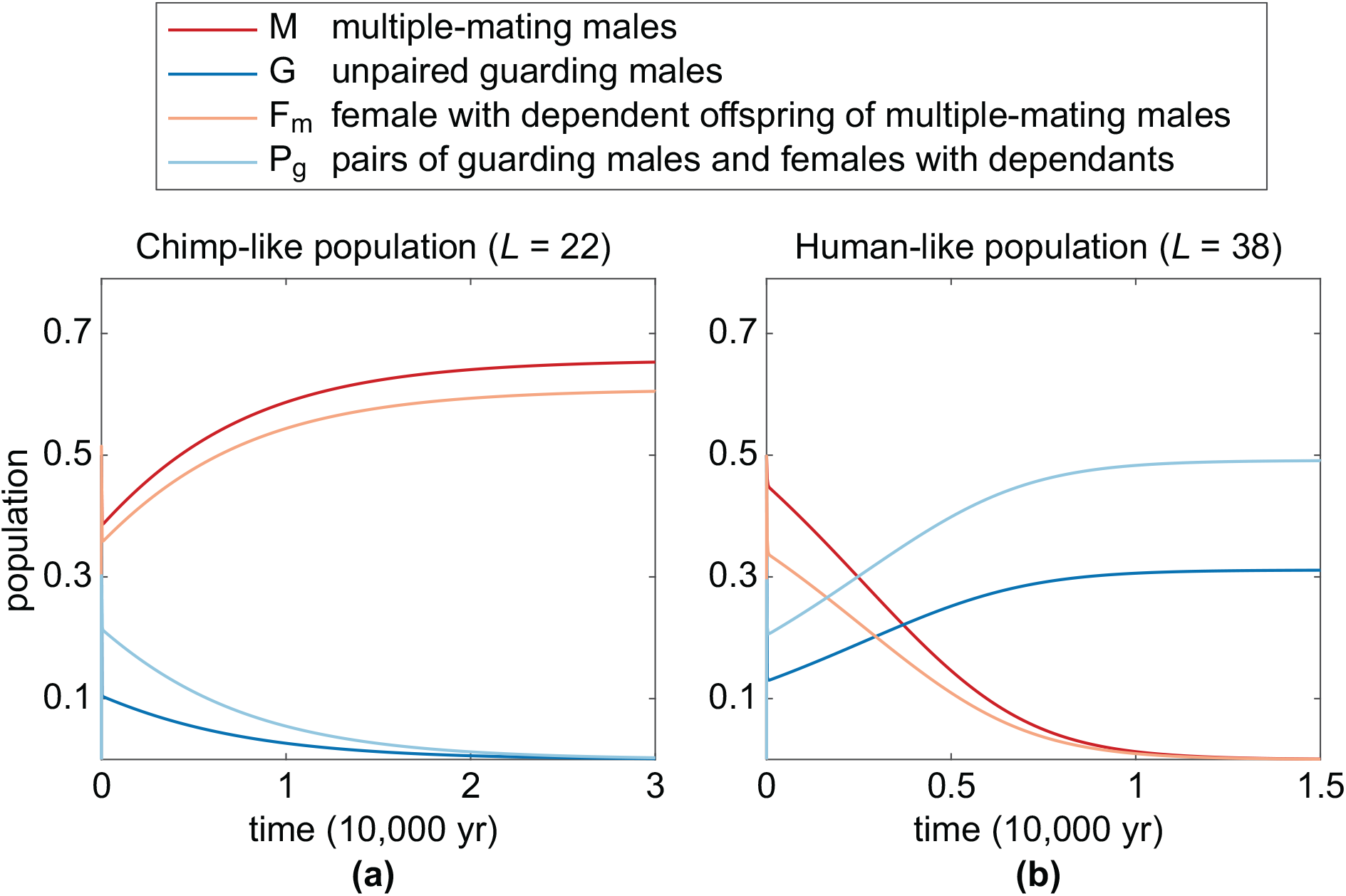
Time evolution of the model for various populations. Solutions are shown with (a) chimp-like and (b) human-like populations. Notice that multiple mating dominates the chimp-like population and guarding dominates the longer-lived populations at equilibrium. All parameters are as in Table 1 with no paternity uncertainty and no pair-bond break up (*q* = 0, *χ* = 0). Initial conditions for all cases are *F* (0) = 1, *M* (0) = 0.5, *G*(0) = 0.5 with all other populations starting at 0.

The model correctly predicts multiple-mating dominates for chimp-like populations and guarding dominates for human-like populations (Figures 5(a) and 5(b)). Note that when mean lifespan increases from *L* = 22 to *L* = 38, there are more unpaired, still-fertile males competing for paternities. In the next section, we analyse the results over parameter space and estimate both the ASR and OSR.

### 3.1 Male-biased sex ratios

We start our analysis by observing the parameter space and the strategy that dominates each point of that space. To do this, we generate a grid of points in parameter space for a range of parameter combinations. In each case, we track males who employ the multiple-mating and mate-guarding strategies. The strategy that results in the largest population at equilibrium is the winner and the corresponding colour is filled in. On each grid, points representing average human- and chimpanzee-like parameters (see Table 1) are identified for comparison. Unless otherwise noted, all of the plots allow for a non-zero rate of fertility ending corresponding to Case 2 in Table 1. Initial conditions were kept the same for each set of parameters: *F* (0) = 1, *M* (0) = 0.5, *G*(0) = 0.5, and all other populations starting at 0. Additionally, both the ASR and OSR (as defined in Table 2) are determined for the model at equilibrium. These ratios are included to illustrate the underlying dynamics of the population and to predict the transition between regions where one strategy dominates.

A comparison of the OSR and ASR when the paternity uncertainty and pair-bond break-up rate are zero is illustrated in Figure 6. Points corresponding to chimp- and human-like populations (Figures 5(a) and 5(b)) are included for comparison. Note that these two points lie on the boundary between regions where the multiple-mating and mate-guarding strategies dominate. Both ratios seem to provide a good indication of where the strategy shift is likely to occur. As both ratios increase, there is greater competition among males for each available female. These larger ratios are caused, in part, by longer interbirth interval lengths. When each offspring requires more years of care, females are out of action for a greater proportion of their fertility window. This effect is offset by a greater expected longevity for which the fertility window is lengthened, which causes the contour lines to slope downward. The same figure for the case without loss of fertility (*ω* = 0) for the same set of parameters is shown in the Supplementary Material. In this case, multiple mating is the only strategy that survives. Note that points corresponding to both chimp- and human-like populations lie along the boundary of the two regions and that they have very similar sex ratios. This is likely due to the simplifying assumptions placed on the model and will be discussed further in the conclusion. Despite this, we can obtain insights into the mechanisms underlying the evolutionary shift in male mating strategies to long-term pair bonds.

**Figure 6:**
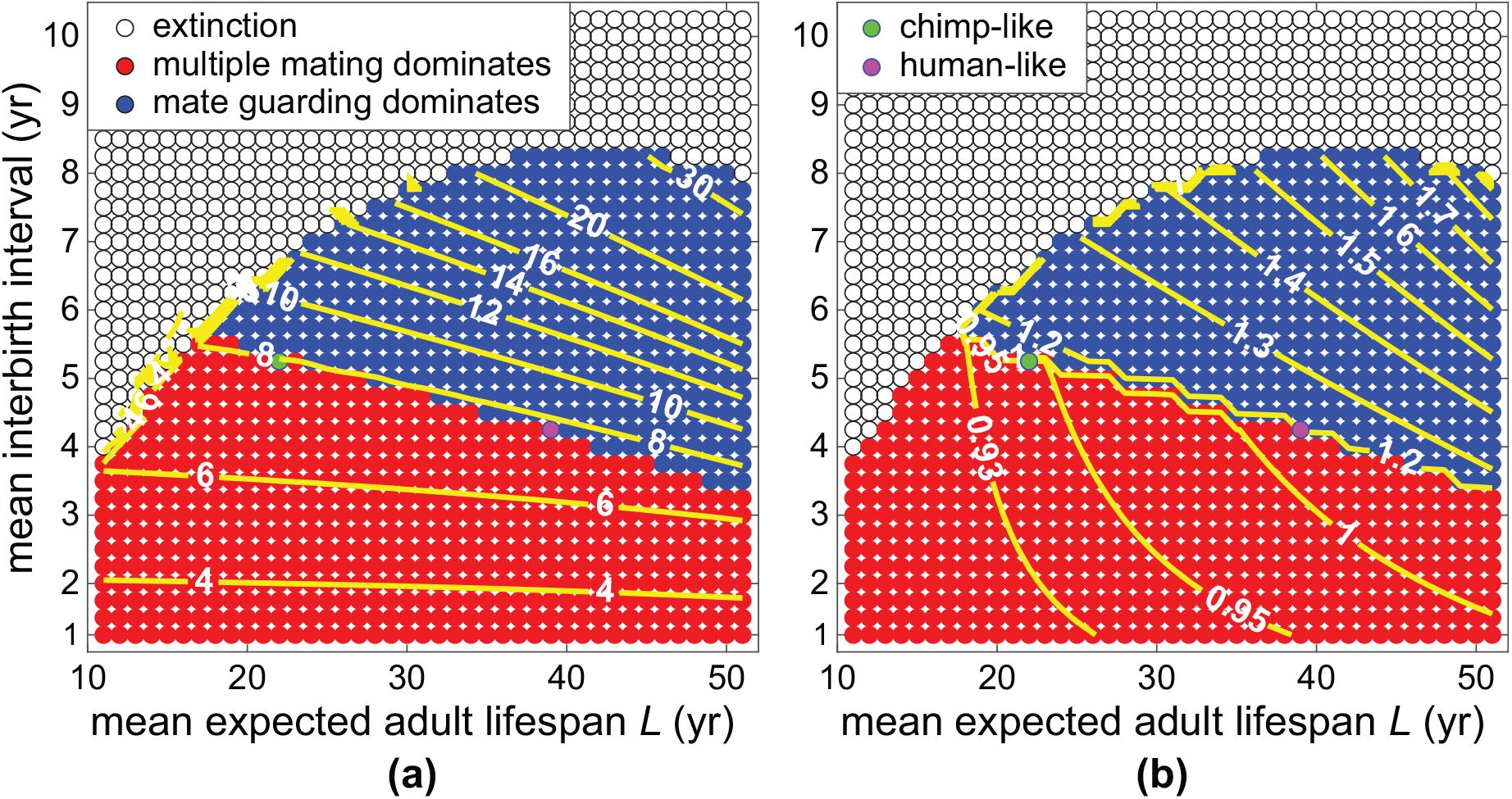
Dominant strategy when interbirth interval and mean expected adult lifespan are varied with contour lines of constant OSR and ASR. (a) The contour lines of constant OSR and (b) ASR seem to provide a good indication of where the strategy shift is likely to take place. Parameters and initial conditions are the same as in Figure 5.

We investigate the effects of increasing paternity uncertainty, *q*, and pair-bond break-up rate, *χ*, in Figure 7. To do this, we start with the baseline model corresponding to Figure 6 and vary *q* and *χ*. On each figure, contour lines of constant OSR and ASR are included that fall closest to the boundary between strategies.

**Figure 7:**
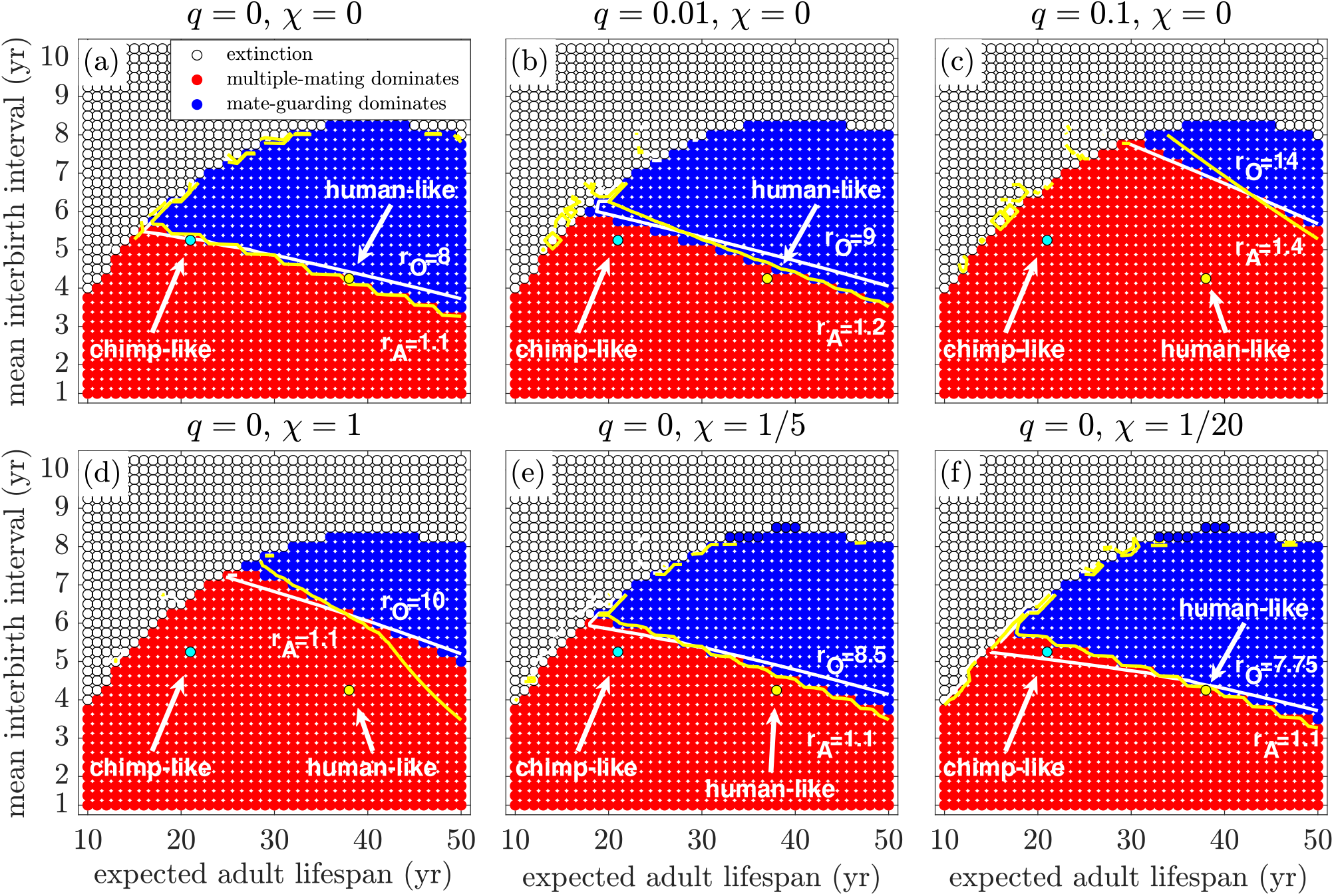
The effect of paternity uncertainty, *q*, and pair-bond break-up rate, *χ*, on the dominant strategy. The baseline model with both paternity uncertainty and pair bond break-up rate set to zero, (a), is compared with varying these parameters. Paternity uncertainty is increased from (b) 1% to (c) 10%, which results in a shrinking guarding region. Additionally, as the pair bond break-up rate decreases from an average duration of (d) 1 year to (e) 5 years and (f) 20 years, it results in a growing guarding region. Points corresponding to chimp and human parameters are indicated with green and yellow circles for comparison (see Table 1).

Figures 7(a), (b) and (c) illustrate the result of increasing the paternity uncertainty, *q*. As expected, greater paternity uncertainty makes it less advantageous to invest effort into mate guarding. If there is a greater chance that multiple-mating males steal the paternity, the advantage of guarding rapidly declines. Thus, as paternity uncertainty increases, the mate-guarding region shrinks, and the human-like population is out of this region by the time uncertainty reaches 1%. This effect is even more pronounced when uncertainty is at 10% (Figure7(c)). An increasing paternity uncertainty means that the sex ratio must be more male biased for there to be sufficient incentive to favour guarding over multiple mating, thus the higher ASR and OSR contour lines.

In Figure 7(d), (e) and (f) we explore how the pair-bond break-up rate affects the system. As the break-up rate drops, the mate-guarding region increases in size. This follows expected male behaviour, since it is less advantageous to invest in mate guarding if the average pair bond is short lived. Note that the OSR and ASR on the boundary of the region drops as pairs last longer. This implies that shorter pair duration requires the sex ratio to be more male biased for there to be sufficient incentive to favour guarding over multiple mating.

Another view that clarifies how these two parameters affect the dominant mating strategy is shown in Figure 8. In this case, we fix a single point in the L-*β* space corresponding to a population between chimps and humans for which *L* = 30 and *β* = 5. We observe that a curve of constant OSR of between 8.3 and 8.5 roughly corresponds to the boundary between the regions where guarding or multiple mating dominate (approximately 8 males for each female). Increasing paternity uncertainty causes guarding incentive to drop as the expected payoff is reduced. In contrast, longer pair bonds cause an increase in guarding incentive, since this guarantees more paternity opportunities in the long run. However, there is a trade-off between these two parameters. If pair-bond duration is big enough, it can overcome a greater degree of paternity uncertainty as there will be more overall paternity opportunities, even if the uncertainty is higher.

**Figure 8:**
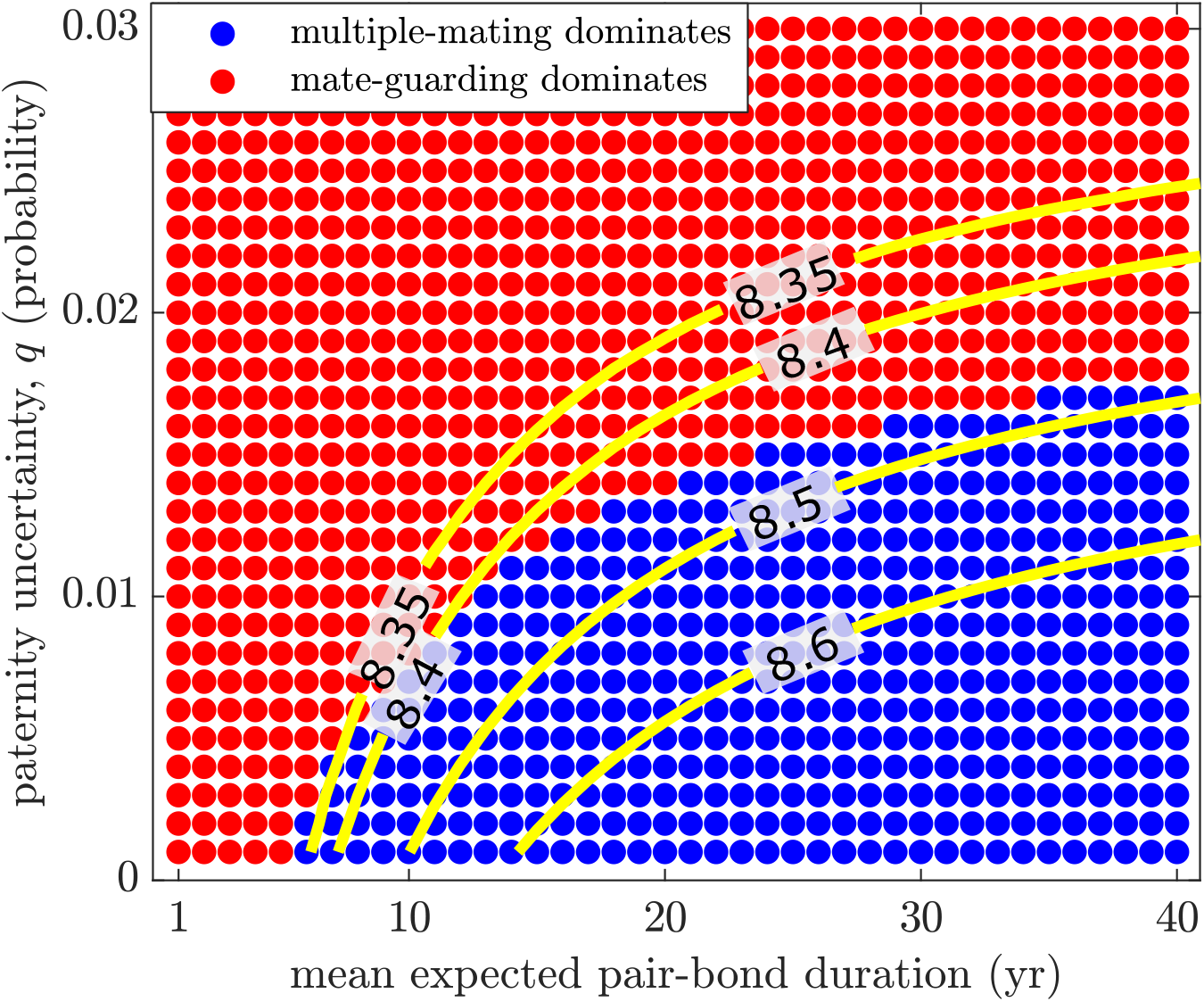
Dominant strategies when paternity uncertainty and mean expected pair-bond duration vary in a population between humans and chimps (*L* = 30, *β* = 5). Included are several lines of constant OSR, which provide an approximate indication of where the strategy shift is likely to take place. Parameters are as in Table 1.

Our model reveals interesting aspects of male mating strategies and shows how the OSR relates to the transition between these strategies. In the next section, we explore the underlying mechanisms that lead to this behavior.

### 3.2 Lifetime paternity opportunities

In this section, we introduce a measure of the average lifetime paternity opportunities (LPO) a typical male following a fixed mating strategy may have. We observe how this can predict the optimal strategy under varying constraints placed on the model. Figure 9(a) demonstrates how the OSR is affected by the interbirth interval for several fixed expected adult longevity values, *L*. Note that as the interbirth interval increases, the OSR displays an exponential increase. This occurs because as the birth interval increases, the population grows slower over time. As a result, there is a higher proportion of older, post-fertile females. Since we assume males are fertile throughout their lives, this implies that the OSR will increase as a consequence. Another view is provided by Figures 9(b) for several fixed interbirth interval lengths, *β*^*−*1^, and varying values of *L*. In a similar way, we see that as longevity increases, the proportion of older, still-fertile males increases, leading to higher sex ratios. The sex ratios measure the competition that each male faces for a single paternity opportunity, so as seen in Figures 6 and 7, increasing sex ratios mean the likelihood a given multiple-mating male will be successful decreases.

**Figure 9:**
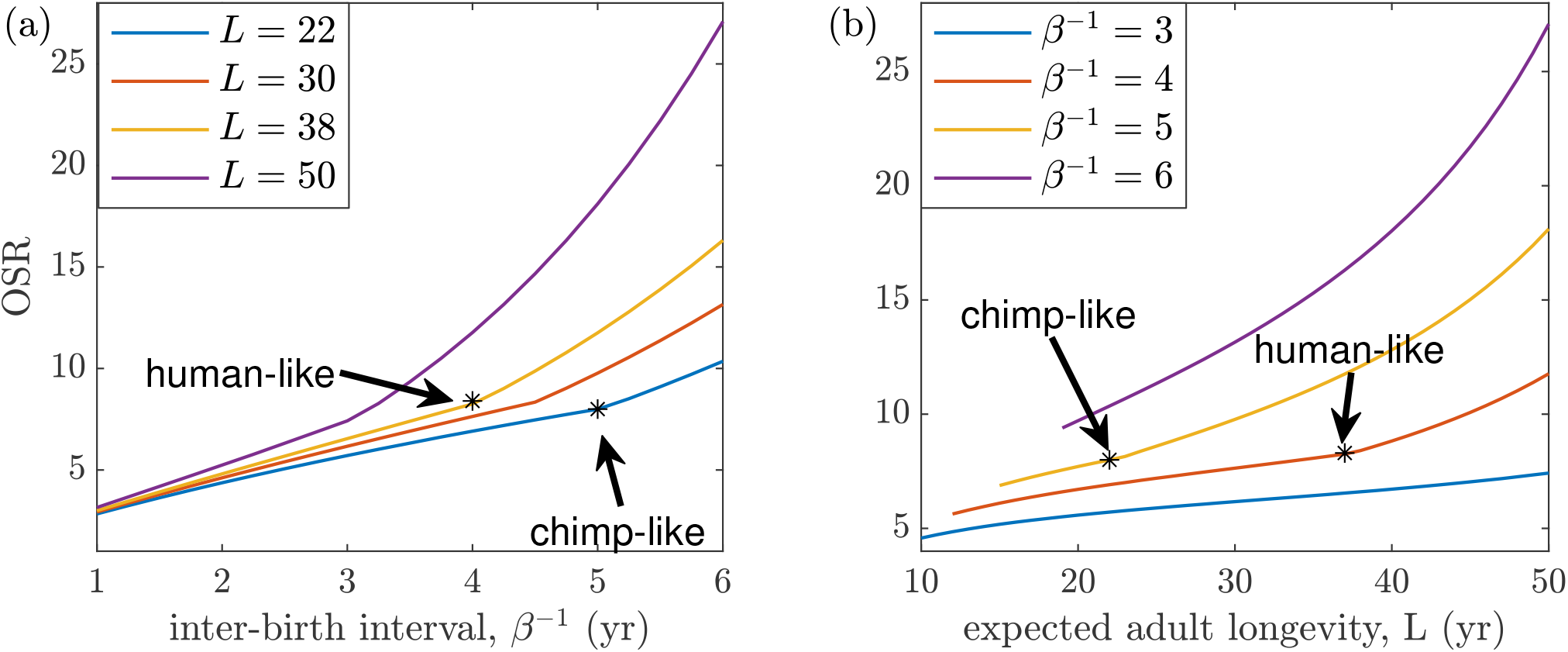
Operational (OSR) sex ratio as a function of interbirth interval and longevity. (a) For several fixed *L* values, the OSR is shown for increasing interbirth intervals. The OSR displays an exponential increase as the interbirth interval increases in length. A lengthening interbirth interval causes the poplulation to grow slower over time. This results in a higher proportion of older, post-fertile females in the population. (b) For several fixed interbirth interval lengths (*β*^*−*1^), the OSR is shown for increasing *L* values. Missing line segments represent populations that reach extinction. As both expected adult longevity and interbirth interval length increase, the OSR responds with an exponential increase. Points corresponding to average chimpanzee and human populations are added for comparison. Other parameters are as in Table 1 with *q* = 0 and *χ* = 0.

The LPO is the expected number of paternities over a male’s lifetime. Given a fixed expected lifespan, *L*, length of offspring dependence, *β*^*−*1^, and operational sex ratio, *r*_*o*_, the maximum number of paternities can be determined for a single male. We compute this quantity separately for males that follow the guarding, Ω_*G*_, and multiple-mating strategies, Ω_*M*_. By observing the relationship between these quantities, we can predict how guarding may have been affected by in increasing OSR. The precise mathematical expressions for Ω_*M*_ and Ω_*G*_ are described in Appendix C.

Figure 10 illustrates the relative advantage for guarding males in both chimp- and human-like populations. The percentage change in the total expected number of lifetime paternities when changing from a multiple-mating, Ω_*M*_, to guarding, Ω_*G*_, is displayed for a range of OSR values. Note that in each case for fixed interbirth interval, *β*^*−*1^, and expected longevity, *L* (chimps: *β*^*−*1^ = 5 and *L* = 22, humans: *β*^*−*1^ = 4 and *L* = 38), a higher OSR results in an advantage for guarding. In particular, note that the model predicts that the OSR for humans and chimps is near 8. At this OSR value, the human-like population can expect an average gain of approximately 68% in total lifetime paternities. This is contrasted with a 31% gain from the same shift in chimp-like populations. Thus, there may not have been enough of an incentive to alter the ancestral mating strategy.

**Figure 10:**
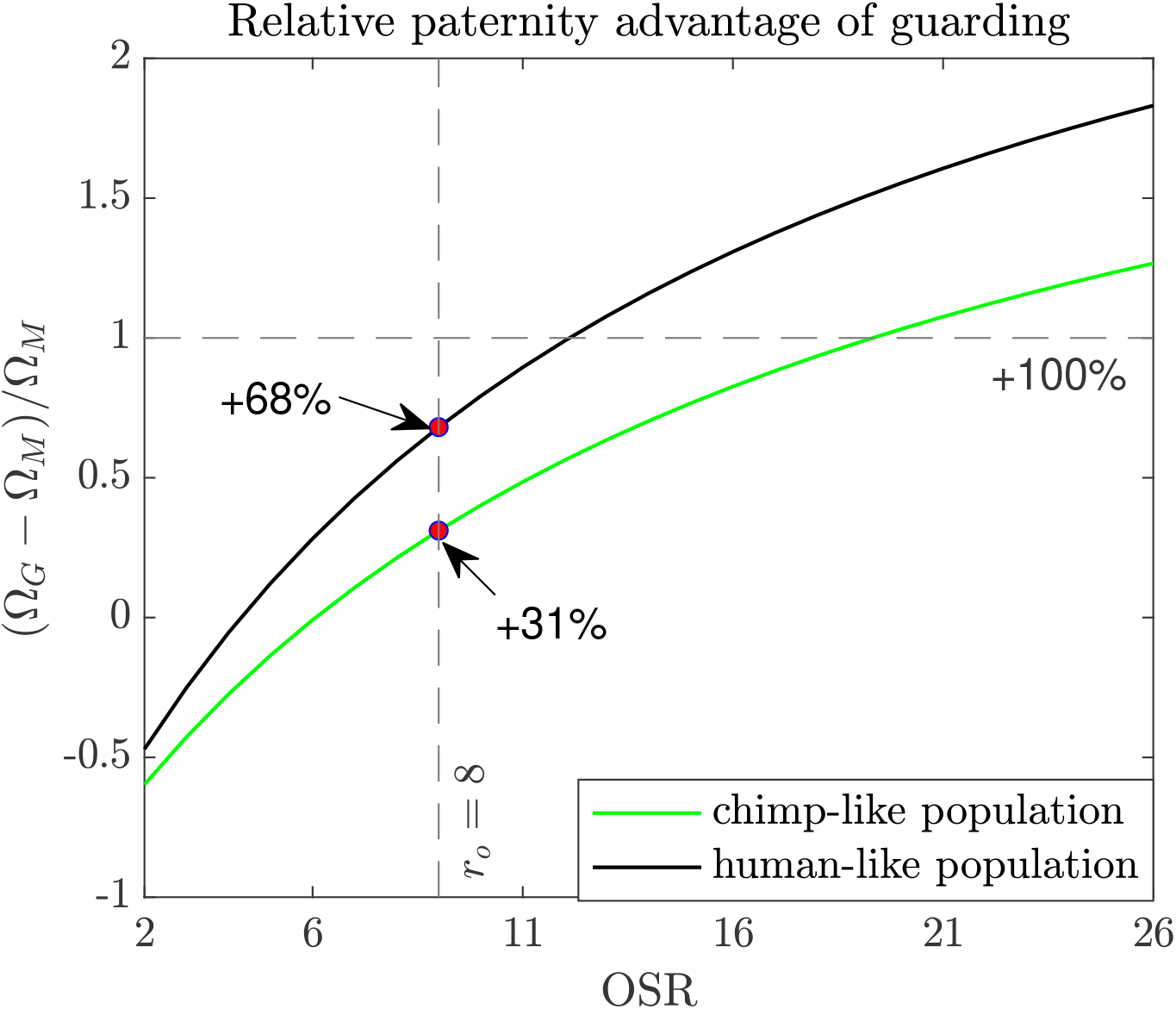
The relative advantage of guarding. The percentage change for changing to a guarding strategy for both human- and chimp-like populations as a function of the OSR are shown. Positive values represent a percentage increase from the ancestral mating strategy. Note that from Figure 6, the OSR for human- and chimp-like populations is approximately 8. At this fixed OSR, there is a bigger advantage for human-like populations. Chimps: *L* = 22, *β*^*−*1^ = 5, Humans: *L* = 38, *β*^*−*1^ = 4.

When the interbirth interval is low, females must provide care for a shorter period of time and are thus free to mate earlier. It also means that paired males do not have long to wait for the next paternity. With a larger population of free females, this results in a lower OSR, which corresponds to less competition for each additional paternity.

On the other hand, when birth intervals are long, paired males must wait longer for the next paternity. It also means that there are fewer free females in the population, resulting in a higher OSR, corresponding to an increased competition for each additional paternity.

This is further complicated by a changing expected adult longevity, *L*. Recall that we approximate the start of female sexual maturity at age *L/*2. Thus, since we assume fertility ends for all females at age 45, the lifetime fertility window for females decreases with increasing *L*. A shorter fertility window means that there are fewer fertile females in the population (recall that we additionally assume that paired males return to the mating pool when the fertility of their partner ends). This is why we observe a slightly larger guarding region with bigger *L*.

However, our model predicts that the bigger effect is the length of the interbirth interval. We observe a tradeoff between these competing forces in the population. Even though males must wait longer for each paternity, the reduction in available females and resulting increase in competition for each paternity gives an advantage to males who guard. This advantage is further strengthened by longer-lived populations as there are even more still-fertile males. This effect is quantified through the LPO which helps to predict the partition of the parameter space into regions where one strategy dominates.

## 4 Conclusion

We have developed a mathematical model that allows us to explore the parameters that contributed to the evolution of human pair bonds. Our model is guided by the argument that connects the emergence of these pair bonds to the male-biased mating sex ratios that accompanied the evolution of human life history [3]. We wish to use this model and subsequent models to help explain the differences between humans and our closest living relatives, the chimpanzees. This model does not assume anything about the details of male-male or male-female interactions, only their final outcomes as they relate to mean reproduction and mortality rates.

The results indicate that scarcity of free, unpaired females relative to the number of unpaired males predicts the likelihood that guarding will be preferred over multiple mating. We’ve observed that this phenomenon is captured in both the ASR and OSR and consequently these ratios can be used to predict a shift in strategy. A more detailed understanding of how expected adult lifespan, interbirth interval lengths, and the male-biased sex ratios combine to affect the strategy that dominates was illustrated through the LPO. This may provide a more accurate index by which to predict the dominant strategy employed by males.

In chimps, the interbirth interval is approximately five years. However, in humans it is on average four years or less, which is surprising given our longevity. The grandmother hypothesis suggests that subsidies from grandmothers to the offspring of their daughters shortened this interval; it was those subsidies that propelled the evolution of increased longevity without favouring continued female fertility to older ages [11, 9, 21, 20]. In our model, this difference, along with greater longevity and longer-lasting pair bonds, allows mate-guarding to remain dominant in humans, which further supports [3]. Nevertheless, this guarding dominance only holds if paternity uncertainty and the pair-bond break-up rate remain low.

Although we obtain intriguing results through analysis of this model, there remain limitations and questions. Considering the differences in human and chimp behaviour, why do points corresponding to these two populations remain so close to the boundaries between mating strategies? In our simulations, it is not obvious that chimps will adopt multiple mating while humans adopt mate guarding. This is likely a consequence of the simplifying assumptions of the model as no age structure is included. This can be further investigated by explicitly considering age structure, e.g., with an agent-based model. Additionally, direct patrilineal inheritance of strategy is unrealistic. It would be interesting to understand how a propensity to adopt a particular strategy is inherited or adapted to social and cultural influences within a lifetime. Also, how does male-male competition affect the outcome? The average age of first reproduction is female-biased (17 for females and 21 for males [26, 17]). Even though the younger males are in the fertile ages, they don’t get paternities because the older males out-compete them. This is a divergence from chimpanzee behaviour where adolescent males actually get paternities on the young females as the older males (who prefer older females [28]) don’t prevent them [27]. Thus, there are important dynamics not captured by this model. Also, there may be other strategies males employ that affect the winning strategy at equilibrium. Guarding males may aid their offspring through provision of care and protection from infanticide, which is not included in this model.

Finally, how did guarding emerge in the first place? There is a long history of debate around how this may have evolved in our lineage and was likely influenced by a range of ecological and cultural factors. The predominant theory is that this behaviour evolved as a consequence of the benefits of cooperative parenting [37, 16, 25, 5, 19, 18]. One of the first models that explored alternative explanations found that there are broad conditions under which guarding became the dominant strategy [12]. Our model provides support for the hypothesis that human pair bonds evolved with increasing payoffs for mate guarding, which resulted from the evolution of our grandmothering life history [3]. However, there are many aspects of this process still left to be explored.

## A Derivation of the density-dependent death rate *μ*(*t*)

In the main text, we use a density-dependent death rate *μ*(*t*), which contains two components:

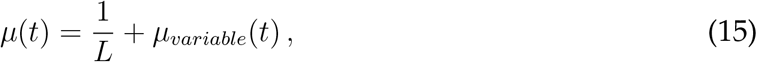

where the first term is a constant death rate given by the reciprocal of the individual’s expected adult life span, *L*, and the second term *μ*_*variable*_(*t*) *≥* 0, is a variable density-dependent death rate. The constant component represents a minimal death rate that ensures each individual will have a maximum average expected adult lifespan of *L* regardless of the variable, density-dependent component.

To determine *μ*(*t*), we first assume *μ*(*t*) is such that the total adult population remains constant, which is possible when the growth rate exceeds the minimal death rate 1*/L*. Note this assumption implies the system has infinitely many equilibria, since any total initial adult population will remain constant. Nonetheless, this definition allows us to compare relative growth rates of the two male mating strategies without introducing new death-rate parameters.

Let *T*_*F*_ = *F* + *F*_*m*_ + *P*_*g*_ + *P* + *F*_*g*_ + *X* + *X*_*m*_ + *X*_*g*_ be the total number of adult females and *T*_*M*_ = *M* + *G* + *P*_*g*_ + *P* + *Y* be the total number of adult males. Since the total adult female and male populations remain constant,

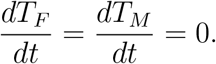

In addition, let us assume we start with an equal number of adult males and females, so that *T*_*M*_ = *T*_*F*_ = *k* for some constant *k >* 0.

We now find the density-dependent death rate *μ*(*t*) in terms of other variables. We have

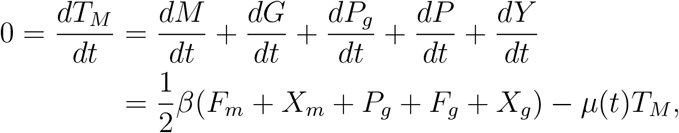

So

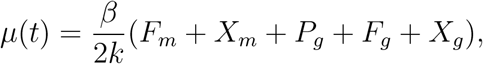

since *T*_*M*_ = *k*. We obtain the same expression if we use *dT*_*F*_ */dt* = 0.

For simplicity, we can normalise all the population variables in (1)–(10) by *k*, so that the total male and female populations *T*_*M*_ = *T*_*F*_ = 1, and we are left with

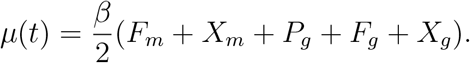

Note we could go further and completely nondimensionalise the system by normalising time *t* by 1*/β*, but since we will numerically simulate solutions, this step hardly reduces computational complexity and the resulting dimensionless parameters would be less convenient, since we intend to vary all life history parameters independently.

Hence, from (12), it follows that

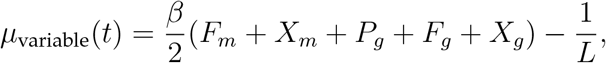

whenever this expression is nonnegative, and *μ*_variable_(*t*) = 0 otherwise, so the expression for our density-dependent death rate is

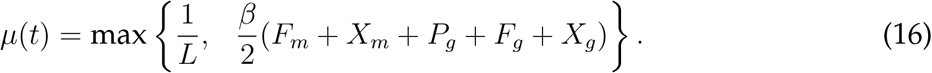

## B The case with no female fertility loss

Figure 11 shows results when there is no end to female fertility. In this case, the multiple-mating strategy always dominates. The adult sex ratio is no longer male-biased, so less competition leads to multiple-mating behaviour. Additionally, notice that the interbirth interval has a smaller effect on extinction when *L* increases. If females are able to give birth throughout their entire lives, longer maturation periods do not lead to extinction.

**Figure 11:**
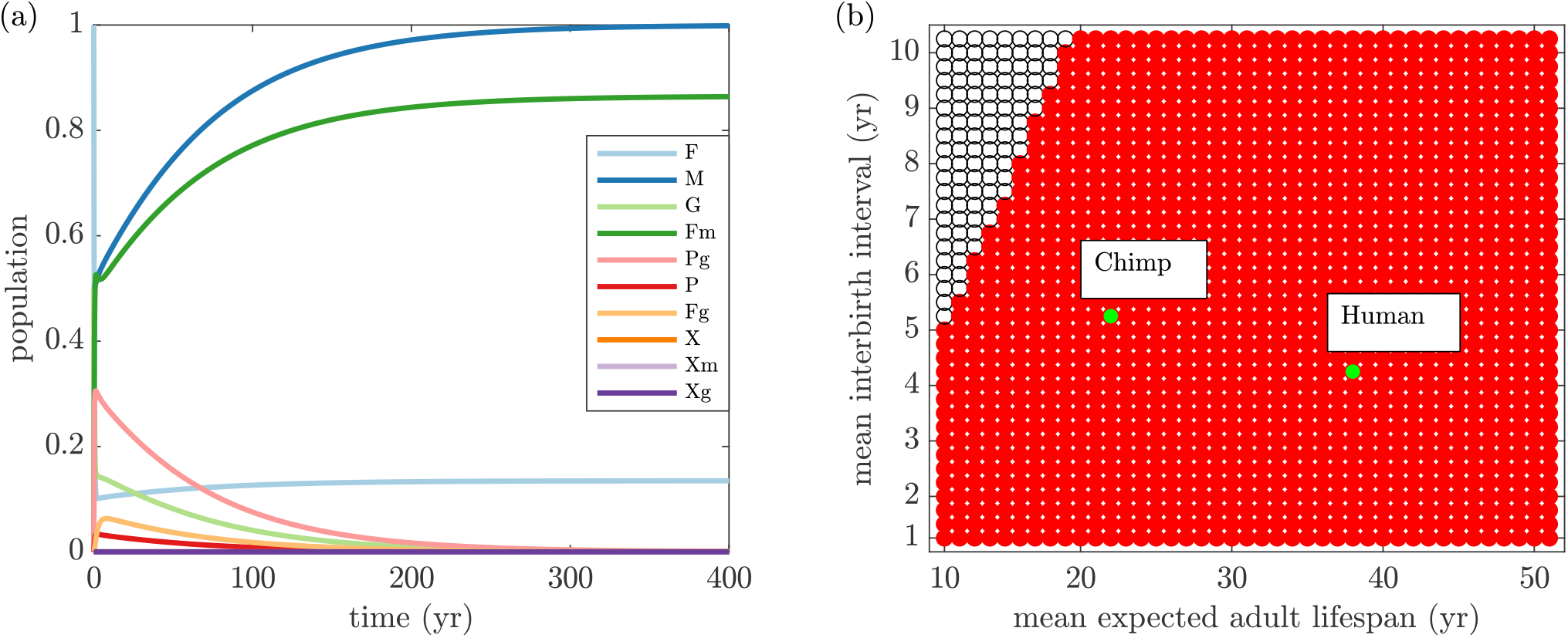
Results when there is no end to the fertility period for females. (a) Trajectories for populations with average adult longevity *L* = 38. In this case, there is no male-biased adult sex ratio and multiple-mating dominates. (b) The dominant strategy is multiple-mating for all values of expected adult lifespan, *L*, and the rate of offspring maturation, *β*. Notice in this case there is far less extinction as both *L* and *β* increase since females can give birth up until death.

## C Calculation of lifetime paternity opportunities

To calculate the lifetime paternity opportunities (LPO) for fixed OSR, *r*_*o*_ = (*M* + *G*)*/F*, we fix the death rate at *μ* = 1*/L*, take the original system (1)–(10)and remove the post-fertile female population without dependants as it will not add to the sum. Thus, we consider the augmented system

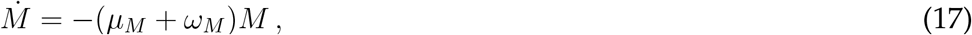

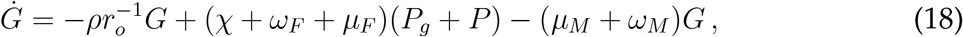

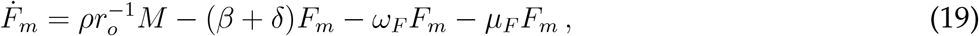

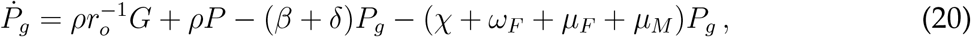

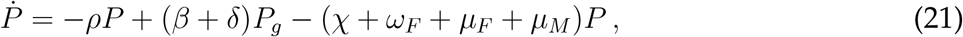

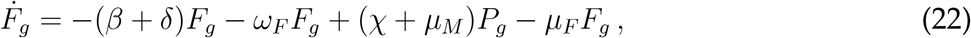

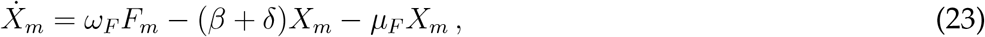

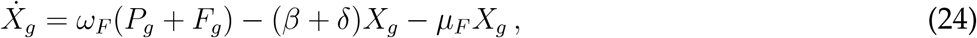

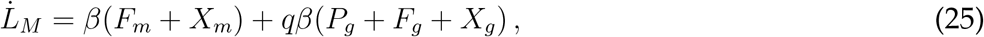

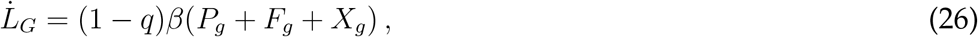

where *L*_*M*_ and *L*_*G*_ represent the cumulative number of paternities for the multiple mating and guarding males at time *t*. Thus, we define the LPOs for multiple mating and guarding males as

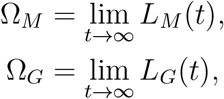

respectively. Starting with initial conditions of *M* (0) = 1 and all other populations equal to zero will give the LPO of multiple-mating males. Similarly, starting with initial conditions of *G*(0) = 1 and all other populations zero will give the LPO of guarding males.

Since (17)–(26) is linear, we can solve it explicitly using Wolfram Mathematica to obtain the general form

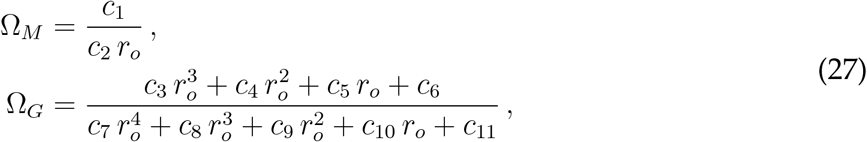

where the *c*_*i*_ = *c*_*i*_(*L, β*), *i ∈ {*1, …, 9*}* are coefficients that are functions of both *L* and *β* (shown in full at the end of this section). Thus, for each point in parameter space, these are constant. In particular, the corresponding LPOs for human- and chimp-like parameters are determined by using the values from the main text (Chimps: *L* = 22, *β* = 1*/*5, Humans: *L* = 38, *β* = 1*/*4)

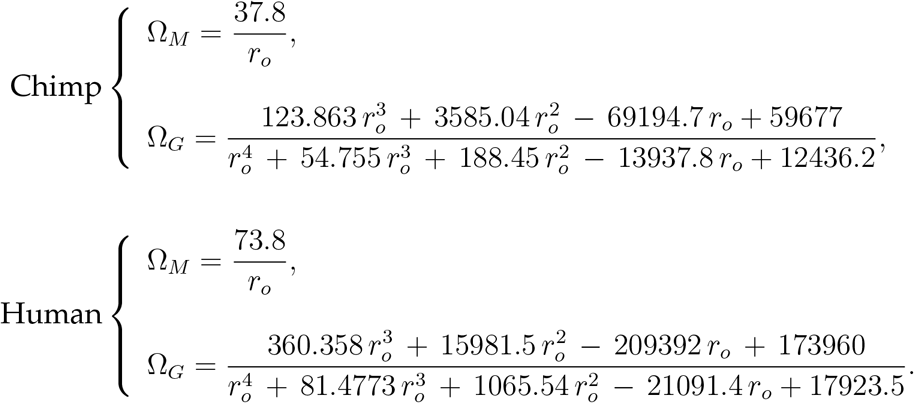

